# Detection of residual disease after neoadjuvant therapy in breast cancer using personalized circulating tumor DNA analysis

**DOI:** 10.1101/425470

**Authors:** Bradon R. McDonald, Tania Contente-Cuomo, Stephen-John Sammut, Ahuva Odenheimer-Bergman, Brenda Ernst, Nieves Perdigones, Suet-Feung Chin, Maria Farooq, Patricia A. Cronin, Karen S. Anderson, Heidi E. Kosiorek, Donald W. Northfelt, Ann E. McCullough, Bhavika K. Patel, Carlos Caldas, Barbara A. Pockaj, Muhammed Murtaza

**Affiliations:** Center for Noninvasive Diagnostics, Translational Genomics Research Institute, Phoenix, AZ, USA.; University of Cambridge, Cambridge, United Kingdom.; Mayo Clinic, Scottsdale, AZ, USA.

## Abstract

Accurate detection of minimal residual disease (MRD) can guide individualized management of early stage cancer patients, but current diagnostic approaches lack adequate sensitivity. Circulating tumor DNA (ctDNA) analysis has shown promise for recurrence monitoring but MRD detection immediately after neoadjuvant therapy or surgical resection has remained challenging. We have developed TARgeted DIgital Sequencing (TARDIS) to simultaneously analyze multiple patient-specific cancer mutations in plasma and improve sensitivity for minute quantities of residual tumor DNA. In 77 reference samples at 0.03%-1% mutant allele fraction (AF), we observed 93.5% sensitivity. Using TARDIS, we analyzed ctDNA in 34 samples from 13 patients with stage II/III breast cancer treated with neoadjuvant therapy. Prior to treatment, we detected ctDNA in 12/12 patients at 0.002%-1.04% AF (0.040% median). After completion of neoadjuvant therapy, we detected ctDNA in 7/8 patients with residual disease observed at surgery and in 1/5 patients with pathological complete response (odds ratio, 18.5, Fisher’s exact p=0.032). These results demonstrate high accuracy for a personalized blood test to detect residual disease after neoadjuvant therapy. With additional clinical validation, TARDIS could identify patients with molecular complete response after neoadjuvant therapy who may be candidates for nonoperative management.

**One Sentence Summary:** A personalized ctDNA test achieves high accuracy for residual disease.

## Introduction

To maximize the rate of cure, cancer patients with early stage disease are often treated aggressively with multiple modalities including pre-operative systemic and radiation therapy, surgery and post-operative therapy. However, this results in overtreatment and adverse effects for some patients who could be cured with less intensive treatment(*1*). In many early stage cancer patients, the benefit of each consecutive modality of treatment is not certain. A treatment monitoring biomarker that can accurately distinguish minimal residual disease (MRD) from cure could enable a new paradigm for individualized management of localized cancers, but this has remained elusive because current diagnostics have inadequate sensitivity. In breast cancer, ~30% patients treated with neoadjuvant therapy achieve pathological Complete Response (pathCR) with no histological evidence of invasive tumor in the resected breast tissue and lymph nodes(*2*). pathCR during neoadjuvant therapy is associated with excellent long-term clinical outcomes (10 year relapse free survival rates: HER2+ 95%, TNBC 86% and ER+HER2- 83%)(*3*). In these patients, surgery provides diagnostic value to confirm pathCR but has not been shown to provide any further therapeutic benefit. An alternative diagnostic test to accurately detect residual disease could obviate the need for surgical resection in these patients, but current tests and imaging do not have adequate sensitivity(*4, 5*).

Recent advances in circulating tumor DNA (ctDNA) analysis have shown promise in monitoring early stage cancer patients but these have primarily focused on recurrence monitoring and lack accuracy for MRD detection at any single time point during treatment(*6–9*). In particular, detection of ctDNA after completion of neoadjuvant therapy has been challenging in patients with breast and rectal cancer, even when residual disease is observed at the time of surgery. Due to limited assay sensitivity, previous studies have found no association between molecular Complete Response (ctDNA clearance from blood) and pathCR(*10, 11*). Detection of low levels of ctDNA in early stage patients is impeded by limited blood volumes obtained in a clinical environment, low concentrations of total cell-free DNA and low fractions of tumor-derived DNA in plasma. As a result, sensitivity and analytical precision of ctDNA tests are often limited due to stochastic sampling variation (Fig. 1A).

**Fig. 1.**
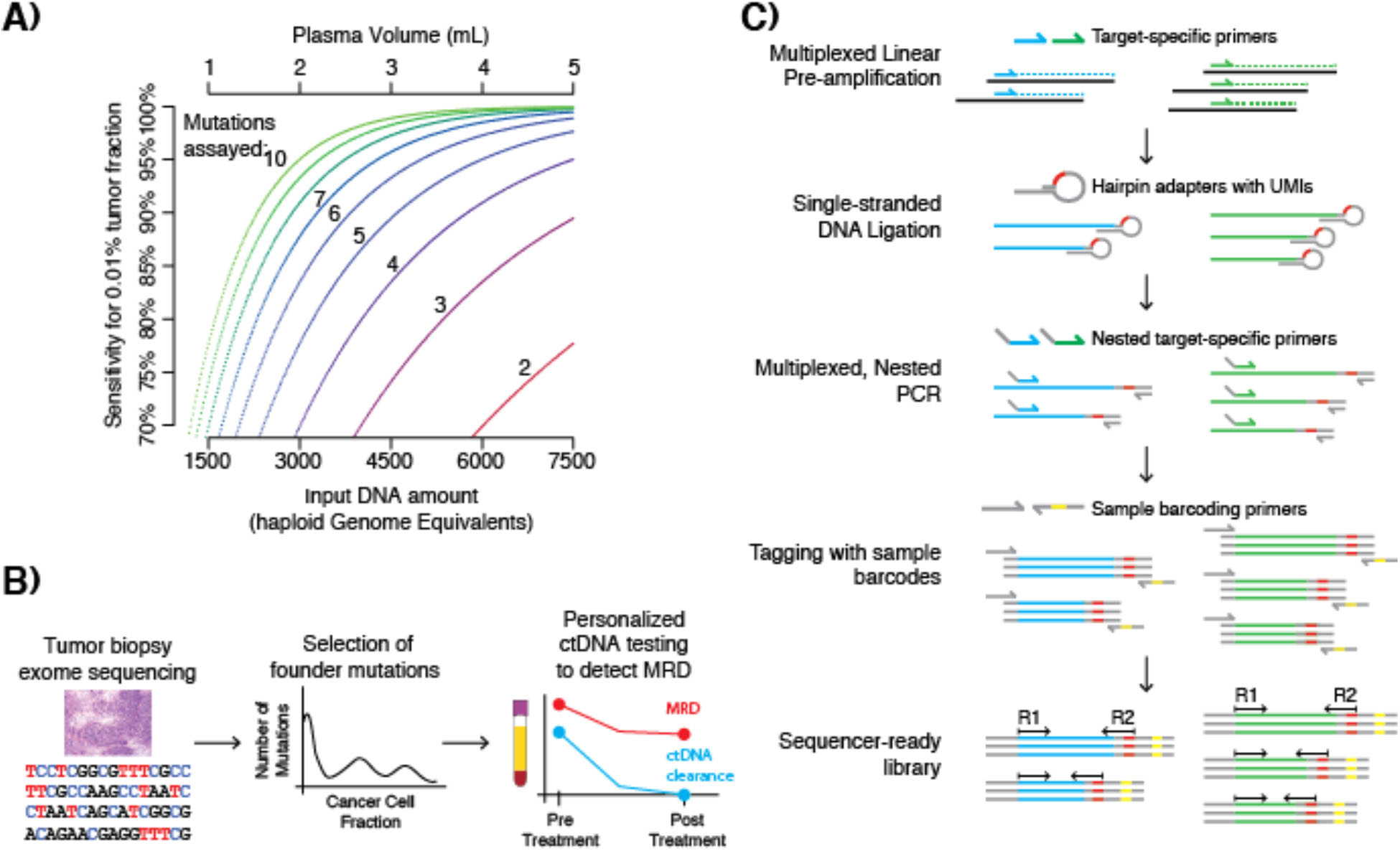
Development of a multiplexed assay for personalized ctDNA detection and monitoring. (A) Based on binomial sampling, maximum theoretical sensitivity for detection of ctDNA at 0.01% tumor fraction is limited if only 1-2 mutations are assayed but can be improved with higher input of plasma DNA and increasing number of patient-specific mutations. (B) TARDIS identifies patient-specific founder mutations using exome sequencing of tumor biopsies and tracks multiple mutations simultaneously in plasma to monitor treatment response and to detect MRD. (C) TARDIS relies on a novel approach for preparing sequencing libraries that includes linear pre-amplification to improve molecular conversion, single-stranded DNA ligation using hairpin oligonucleotides to allow error suppression using template fragment sizes and unique molecular identifiers (UMIs) and multiplexed PCR to enrich targeted genomic loci.

Sampling variation can be overcome by increasing the volume of blood obtained at each time point to increase the amount of total plasma DNA analyzed, by improving the rate of conversion of DNA into sequencing-ready molecules or by simultaneously analyzing multiple patient-specific somatic founder mutations, as these are present in all cancer cells and each is equally informative of tumor-derived DNA in blood(*12*). To leverage these principles and enable MRD detection, we have developed a personalized approach for tumor-guided ctDNA detection and quantification called TARgeted DIgital Sequencing (TARDIS). We identify founder somatic mutations using exome sequencing of tumor biopsies and analyze up to 20 mutations simultaneously in serial plasma DNA samples obtained during treatment (Fig. 1B). To maximize capture and analysis of input DNA while preserving specificity, we perform targeted linear pre-amplification, followed by single-stranded DNA ligation with unique molecular identifiers (UMIs), targeted nested PCR and sequencing (Fig. 1C). The resulting sequencing reads at each targeted locus have a fixed amplification end and a variable ligation end, preserving fragment size information unlike conventional PCR amplicons(*13, 14*). We utilize fragment sizes and UMIs to group sequencing reads into read families (RFs) and require consensus of all members to distinguish true low abundance mutations from polymerase or sequencing background errors (Fig. S1, Supplementary Materials).

## Results

### Evaluation of assay performance in reference samples

To evaluate analytical performance of TARDIS at low ctDNA levels, we designed a multiplexed panel targeting 8 mutations in commercially available reference samples for cell-free DNA analysis (Table S1). We analyzed a total of 93 replicates, 7-16 each at 1%, 0.5%, 0.25%, 0.125%, 0.063%, 0.031% Allele Fractions (AFs) and 16 wild-type (WT) samples. AFs for individual mutations were verified by droplet digital PCR (ddPCR) by the vendor (except for 0.063% and 0.031% that were dilutions of 0.125% in WT, Table S2). Input DNA in each replicate was 5.6-7.9ng (1682-2394 haploid genomic equivalents). Mean number of mutated molecules expected for each targeted mutation in a sample was 0.90-19.6 across 0.031%-1% AFs.

To exclude polymerase errors introduced during linear or exponential amplification, we required at least two independent DNA fragments (≥ 2 RFs) and measured AF consistent with ≥ 0.5 mutant molecules to support each variant call. Using this approach, we observed a mean background error rate of 3.29 × 10^−5^ (Fig. S2, Supplementary Materials). In reference samples, we achieved mutation-level sensitivity of 94.6%, 90.6%, 65.6%, 50.8%, 25.8% and 19.6% respectively at 1%, 0.5%, 0.25%, 0.125%, 0.063% and 0.031% AFs, consistent with decreasing number of mutant molecules at lower AFs (Fig. 2A, Data S1). Using the same criteria, none of the 128 candidate mutations were detected in wild-type samples (100% specificity). Analogous to the detection of tumor-derived DNA in plasma, we leveraged multiple mutations to evaluate sample-level sensitivity. To ascertain mutant DNA in a sample, we required ≥ 2 RFs contributed by one or more mutations, each with measured AF consistent with ≥ 0.5 mutant molecules in input DNA. In samples where a single mutation was detected, we required supporting RFs with ≥ 2 fragment sizes (Supplementary Materials). We achieved sample-level sensitivity of 100% for 0.125%-1% AFs, 87.5% for 0.063% and 78.6% for 0.031% AF (Fig. 2B). Using the same criteria, we detected mutant DNA in 1 of 16 wild-type samples (93.8% specificity). These results demonstrate the principle underlying TARDIS i.e. leveraging multiple patient-specific mutations to overcome limits of sampling and to improve limit of detection. We successfully detected mutant DNA in 11 of 14 replicates with 0.031% AF with 7.8 ng input DNA per reaction, when we expected a total of 7.2 mutant molecules per reaction across 8 mutations (<1 mutant molecule per mutation).

**Fig. 2.**
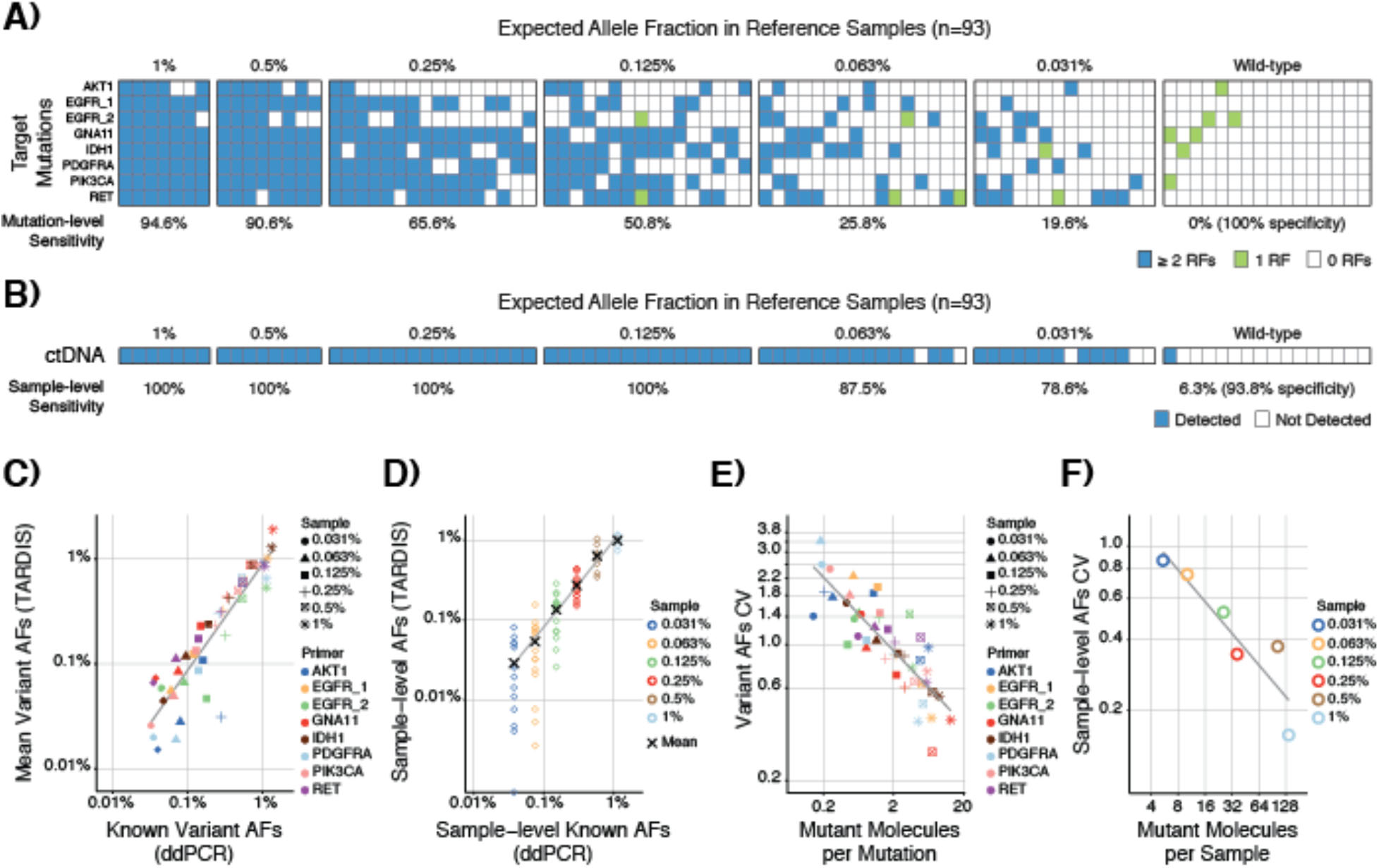
Analytical validation of TARDIS using reference samples. (A) Mutation-level sensitivity and specificity across 93 reference samples and 8 mutations, requiring each mutation is supported by ≥ 2 RFs and an AF consistent with ≥ 0.5 mutant molecules. (B) Sample-level sensitivity and specificity, requiring ≥ 2 RFs contributed by one or more mutations, each with an AF consistent with ≥ 0.5 mutant molecules. (C) Comparison of variant AFs observed using TARDIS (mean for each variant across all replicates at the same mutation level, 48 data points) with known variant AFs measured using ddPCR. Gray line is linear fit. (D) Comparison of sample AFs observed using TARDIS (mean for all 8 mutations assayed in each replicate sample, 77 data points) with known sample AFs (mean of known variant AFs). Gray line is linear fit to the mean at each AF level. (E) CVs of variant AFs decreased with increasing number of mutant molecules per mutation. CVs calculated across 7-16 replicates at each mutation level for each of 8 mutations (48 data points). (F) CVs of sample-level AFs were lower than those for individual mutations, demonstrating the advantage of leveraging multiple mutations for ctDNA quantification. CVs calculated across 7-16 replicates for sample-level means across 6 mutation levels.

To determine quantitative accuracy, we compared known AFs for variants measured by ddPCR in reference samples to mean AFs measured using TARDIS and found a strong correlation (Pearson r=0.921, p<2.2×10^−16^, Fig. 2C). To evaluate agreement between observed and expected mutant fraction in each sample (equivalent to ctDNA fraction in plasma samples), we calculated sample-level mean AFs (mean of 8 mutations in each replicate) and found an excellent correlation between observed and expected AFs (Pearson r=0.937, p<2.2×10^−16^, Fig. 2D). To evaluate quantitative precision, we calculated coefficient of variation (CV) for mutation AFs across replicates and expected AF levels. Across 8 variants at 6 different AF levels, each evaluated in 7-16 replicates, we found CVs agreed with inverse square root of the number of rare mutant molecules (1/√n), consistent with expectations of the Poisson distribution (Fig. 2E). CVs for mutation AFs ranged from 0.28 (17.8 average mutant molecules per mutation) to 3.74 (0.08 mutant molecules per mutation). To evaluate whether multiple mutation analysis improves precision in mutant fraction estimates, we calculated CVs for sample-level mean AFs across replicates and CVs ranged from 0.16 for 1% expected AF (137.9 average mutant molecules per reaction) to 0.87 for 0.031% expected AF (5.4 mutant molecules per reaction, Fig 2F). These results confirm quantitative accuracy of TARDIS and demonstrate that simultaneously assaying multiple patient-specific mutations improves quantitative precision.

### Detection of residual disease in patients with early stage breast cancer

To evaluate whether TARDIS enables MRD detection in early stage cancer patients, we analyzed blood samples obtained from 13 patients with breast cancer treated with neoadjuvant therapy (NAT). We performed whole exome sequencing of DNA from diagnostic tumor biopsies and matched germline samples, achieving 193x and 148x mean coverage respectively (Table S3). We identified and designed primers for 13-150 founder mutations per patient (mean 65.9). By using an aggressive filtering strategy, we retained only high-quality loci with primers predicted to perform adequately in multiplex (9-24 mutations per patient, mean 13.2, Supplementary Materials). After further excluding primers amplifying erroneously in control samples, we analyzed 34 serial plasma samples obtained from 13 patients (2-4 samples per patient, Data S2) for 8-23 mutations per patient (mean 12.2, Fig. 3A, Data S3). Samples were collected prior to commencement of therapy, during NAT and after completion of NAT before surgery. Input plasma DNA amounts were 5.6-34.5 ng per sample (mean 18.0, median 17.2), obtained from 0.2-4.2 mL plasma (mean 2.7, median 3.0) and analyzed in 1-2 replicate TARDIS experiments. We detected ctDNA in 12/12 patients prior to treatment at 0.0019%-1.04% (mean 0.26%, median 0.040%), supported by 2-23 distinct mutation events (mean 7.7, median 6.5, Supplementary Materials) and 4-591 mutant RFs (mean 126.4, median 27.5, Data S4). Baseline plasma sequencing failed in one patient (E009). In 6/13 patients, invasive residual disease was observed at the time of surgery after completion of NAT while 2/13 had in situ residual disease. In 5/6 patients with invasive residual disease, we detected ctDNA in blood after NAT (Fig. 3B). In the sixth patient (T065), ctDNA was undetectable in the last blood sample after completion of NAT, likely due to limited plasma DNA available for analysis (8.7 ng compared to median DNA input of 17.2 ng for all samples analyzed and 23.2 ng for samples obtained after NAT). ctDNA was detected in 2 prior blood samples from the same case obtained 6 and 12 weeks earlier (19.6 ng and 15.0 ng input respectively). In 2/2 patients with in situ residual disease (ypTis N0), we detected ctDNA after completion of NAT. Five patients achieved pathological Complete Response (pathCR) with no evidence of residual tumor. In 4/5 patients with pathCR, ctDNA was undetectable in blood after NAT (Fig. 3B). In the fifth patient with pathCR (T014), ctDNA signal continued to be robustly detected in 3 samples collected at 6-weekly intervals throughout neoadjuvant therapy. Since no residual disease was observed in the surgical specimen, persistent ctDNA detection is suggestive of distant micro-metastasis. In patients with detectable ctDNA after NAT, tumor fraction was 0.0064%-0.046% (mean 0.019%, median 0.017%), supported by 2-6 distinct mutation events (mean 4.0, median 4.0) and 2-21 mutant RFs (mean 12.4, median 15). We observed a decrease in ctDNA levels after NAT compared to pre-treatment levels in all but 2 patients (0.04% median AF at baseline compared to 0.02% median AF after NAT, Wilcoxon signed rank test p=0.0024, Fig. 3C). Temporal changes in variant AFs for multiple mutations within each patient agreed with each other, until affected by sampling variation as ctDNA levels decreased during treatment (Fig. 3D and Fig. S3).

**Fig. 3.**
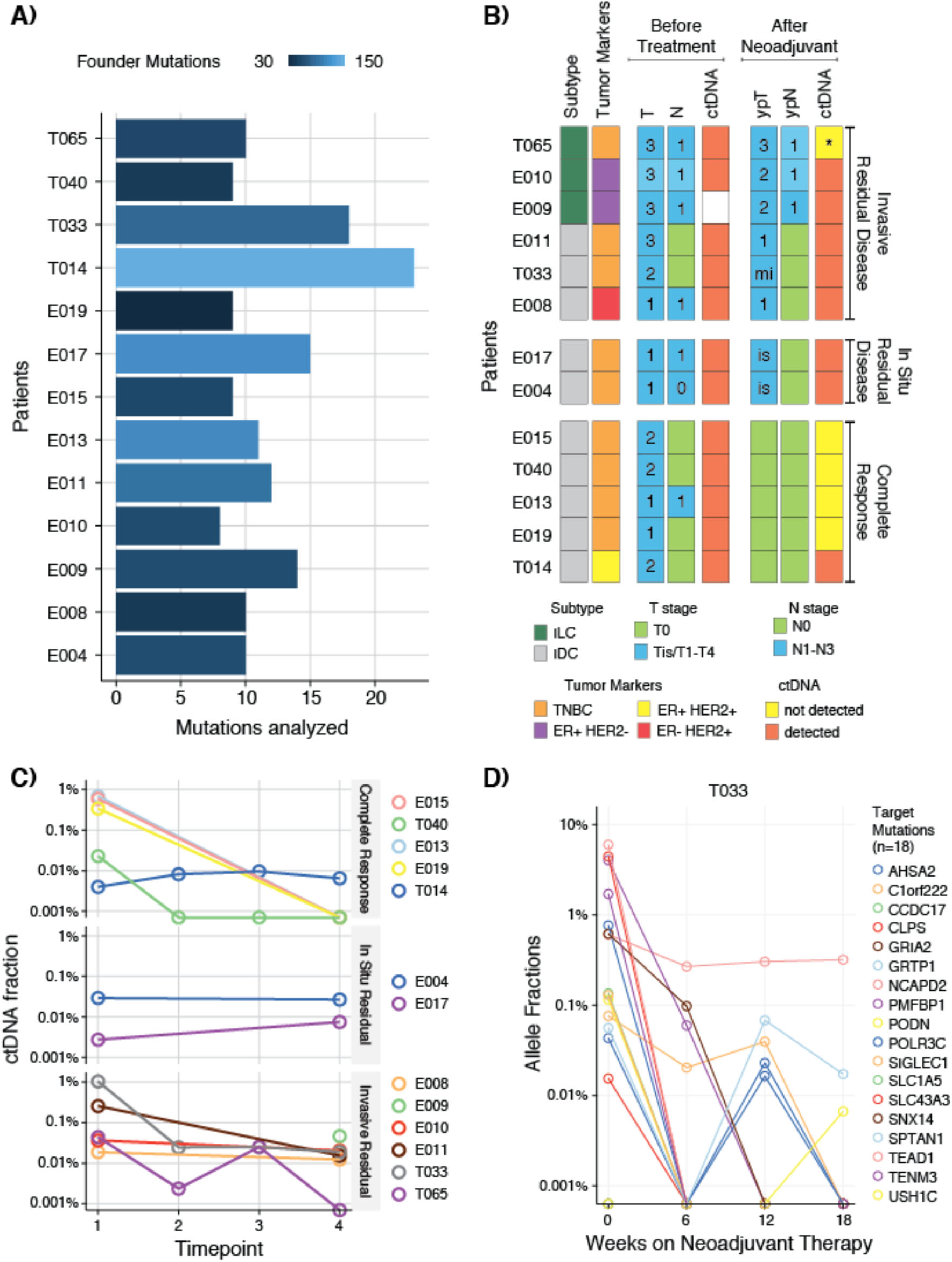
MRD detection in patients with breast cancer following neoadjuvant therapy. (A) Number of mutations analyzed using TARDIS in each patient. Bar colors indicate the number of founder mutations identified using tumor exome sequencing. (B) Summary of clinical data, TNM staging and ctDNA detection before treatment and after neoadjuvant therapy. Pathological TNM staging was performed after surgery and completion of NAT. Number in each box indicates T or N stage, is: in situ, mi: microinvasive disease. *ctDNA was undetectable in the last blood sample collected after completion of NAT for patient T065 but detected consistently in 2 prior blood samples collected 6 weeks and 12 weeks earlier. The baseline plasma sample for E009 failed sequencing. (C) Changes in ctDNA levels during neoadjuvant therapy, grouped by clinical response to treatment (Complete Response, In Situ Residual Disease and Invasive Residual Disease). Time point 1 is before treatment and time point 4 is after completion of NAT for all patients. For T014, T033, T040 and T065, time points 2 and 3 were collected at 6 and 12 weeks during NAT (D) A representative example of changes in individual mutation AFs during neoadjuvant therapy in patient T033. Individual AFs for all patients are shown in Fig. S3.

To evaluate how sensitivity for ctDNA detection was affected by increasing the number of mutations assayed, we sub-sampled combinations of different number of mutations in our dataset (Fig. 4). Using any one of the assayed mutations from each patient, we found mean sensitivity of 37% at baseline and 14.4% after completion of neoadjuvant therapy. Sensitivity for baseline ctDNA detection peaked at 10 mutations and plateaued thereafter. In contrast, sensitivity for ctDNA detection after NAT peaked at 14 mutations.

**Fig. 4.**
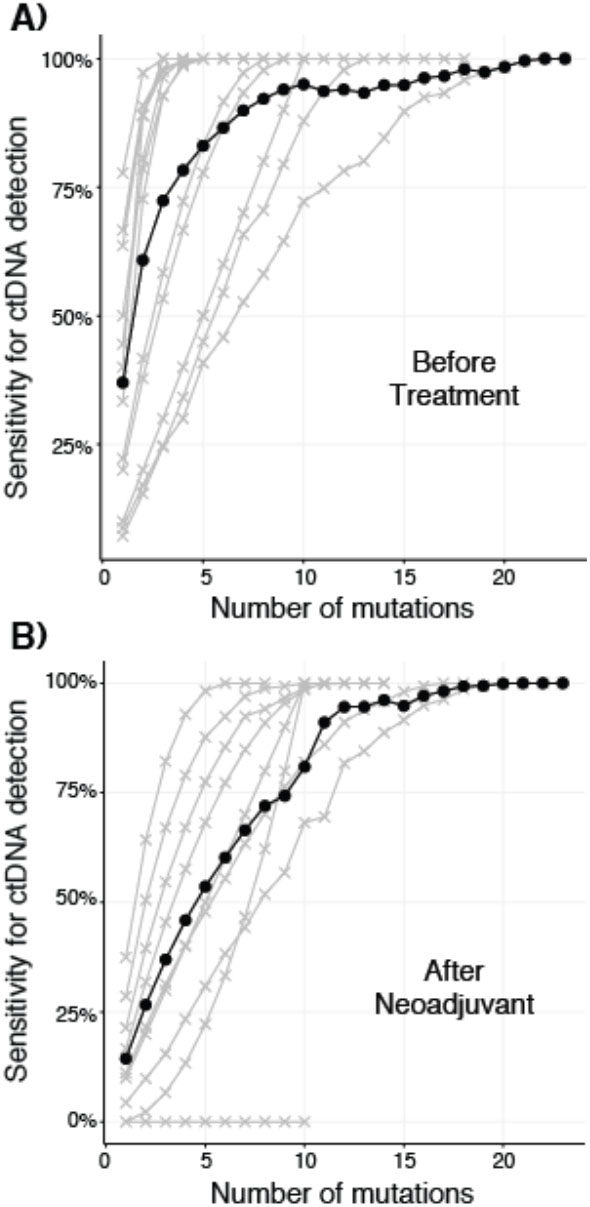
Sensitivity for ctDNA detection improved with increasing number of mutations analyzed using TARDIS, both before treatment (A) and after neoadjuvant therapy (B). Sensitivity for each patient is shown as gray lines with crosses. Mean sensitivity for each number of mutations is shown as black line with dots. For each number of mutations, we sampled combinations from targeted mutations in each patient and tested up to 1000 randomly chosen combinations. Pre-treatment samples from 12 patients are included in (A). Samples obtained after completion of neoadjuvant therapy from 9 patients are included in (B), excluding patients with ctDNA clearance and Complete Response.

## Discussion

Our results demonstrate that ctDNA detection at the end of neoadjuvant therapy for breast cancer is associated with presence of residual disease at the time of surgery (Fisher’s Exact p=0.032, Odds Ratio 18.5). Previous studies have reported ctDNA detection rate of 50-70% before treatment in localized breast cancer patients(*7, 15*). In contrast, we detected ctDNA in all successfully analyzed patients at presentation (significantly higher than published detection rates, p=0.0254). In two recent studies, ctDNA was detectable after completion of neoadjuvant therapy in only 1/22 breast cancer patients (4.5%) and 11/91 rectal cancer patients (12%) when residual disease was found at surgery and no association between ctDNA clearance and pathological complete response was observed(*10, 11*). These studies targeted a single mutation for plasma DNA analysis using digital PCR or digital sequencing and their results are consistent with ctDNA detection rates we observed when evaluating any one of the targeted mutations in our study (Fig. 4B). Using TARDIS to analyze multiple patient-specific mutations simultaneously, we detected ctDNA in 7/8 breast cancer patients with residual disease (significantly higher than previously reported detection rates, p<2.2×10^−16^). Our results suggest that tumor-guided personalized ctDNA analysis using TARDIS is a reliable approach to identify patients with molecular complete response (ctDNA clearance from blood) after neoadjuvant therapy. Together with imaging and tissue-based predictive biomarkers, molecular complete response could become a useful adjunct diagnostic test to individualize decisions about additional treatment such as surgery or adjuvant therapy.

Recent studies have evaluated ctDNA analysis for MRD detection in early stage cancer patients using digital PCR(*7*), amplicon sequencing(*8, 9*) or hybrid-capture gene panel sequencing(*6*). However, reported methods lack adequate sensitivity to reliably detect residual disease at any single time point. Rather, these efforts focused on using ctDNA for monitoring of early relapse during long-term follow-up, showing evidence of ctDNA detection prior to clinically detectable recurrence. Any test that seeks to stratify patients for further treatment after neoadjuvant therapy, including surgical resection and adjuvant therapy, must have an adequate limit of detection for MRD immediately after neoadjuvant therapy or after surgery. TARDIS makes key technical advances that enable MRD detection by selecting and prioritizing founder mutations, simultaneously analyzing up to 20 mutations in multiplex, using linear pre-amplification to maximize the fraction of limited plasma DNA amounts effectively analyzed and suppressing errors by using both, molecular identifiers and fragment sizes. Due to additional sensitivity and specificity provided by these improvements, we reliably detected residual disease after neoadjuvant therapy for breast cancer, detecting tumor fractions as low as 0.002% using 3 mL plasma samples. Recent reports propose genome-wide sequencing or enrichment of patient-specific mutations by hybridization to detect very low levels of ctDNA in plasma(*16–18*). These approaches are currently expensive either due to amount of sequencing data required for adequate genome-wide coverage for each plasma sample or due to the cost of synthesizing and optimizing biotinylated oligonucleotides for enrichment by hybridization for each patient-specific assay. In contrast, TARDIS achieves adequate sensitivity for MRD using cost-effective personalized assays that only require conventional primer synthesis and a limited sequencing footprint. In addition, we prioritize clonal founder mutations for ctDNA analysis that should be present in all cancer cells and therefore, they are unlikely to be lost due to population bottlenecks during treatment and remain informative for MRD detection(*9, 12*). These features enable more frequent and longitudinal analysis of plasma samples once a patient-specific TARDIS panel has been developed.

The limit of detection required to detect MRD using a blood test will vary across cancer types and disease stages. We observed that limit of detection can be improved by increasing the number of founder mutations targeted (Fig. 4). A current limitation of TARDIS is the ability to target ~10-20 mutations in multiplex, imposed by stringent informatic filtering of primers during personalized assay design to minimize off-target amplification. Additional performance data from multiple patient-specific panels will allow us to refine primer design and increase the number of multiplexed target-specific primers. In addition, inclusion of other somatic genomic alternations such as indels and fusion breakpoints can expand the pool of eligible patient-specific target mutations.

Despite a limited clinical sample size, our results provide proof-of-concept for using personalized multi-mutation ctDNA monitoring to predict residual disease after neoadjuvant therapy, an application of ctDNA analysis that has not been previously demonstrated successfully. We did not detect ctDNA in one patient with residual disease, most likely due to a combination of low plasma DNA concentration in the post-neoadjuvant therapy sample and a low rate of ctDNA shedding from the tumor (suggested by low pre-treatment ctDNA levels for a T3N1 tumor). ctDNA was detected in this patient in 2 plasma samples collected 6 weeks and 12 weeks earlier, suggesting that future clinical studies to evaluate MRD detection after neoadjuvant therapy will benefit from analysis of larger blood volumes. Although in the current study, we analyzed up to 4 mL plasma obtained from 10 mL blood samples, it is conceivable to collect up to 30 mL blood at a single time point. It is also feasible to collect and analyze plasma samples over multiple days after completion of neoadjuvant therapy. If ctDNA is cleared from blood and remains undetectable in multiple consecutive samples, this could accurately rule out MRD. Interestingly, we also detected persistent ctDNA signal in one patient with pathCR and no residual tumor, suggesting our approach may be able to detect distant micro-metastasis. This patient with ER+ HER2+ cancer was treated with additional adjuvant systemic therapy as part of routine clinical practice and has not shown any evidence of relapse during nearly 5 years of follow-up. Finally, we also report extensive analytical validation data using commercially available reference samples that could enable benchmarking of current and future technologies for ctDNA analysis.

Overtreatment of early stage cancer patients remains a challenge in cancer medicine, likely to become more relevant as newer blood- and imaging-based early detection approaches gain credence(*19*). Most efforts to optimize treatments have focused on tissue-based predictive biomarkers to assess risk of tumor recurrence(*20*). Our results suggest blood-based MRD testing during treatment can further help evaluate additional benefit of each treatment modality in individual patients. Establishing clinical validity and utility for pre- and post-operative MRD detection will require larger and prospective studies with long-term clinical follow-up. Once validated, using MRD detection to individualize cancer management could substantially reduce treatment-related morbidity while preserving clinical outcomes.

## Materials and Methods

### Patients and samples

This study includes patients prospectively enrolled at Mayo Clinic, Phoenix, AZ, USA under an approved IRB protocol number 14-006021 (Mayo cohort) or at Addenbrooke’s Hospital, Cambridge, UK under an approved Research Ethics Committee protocol number 12/EE/0484 (Cambridge cohort). Informed consent was obtained from all patients. Tumor samples obtained at the time of diagnosis were exome sequenced. Blood samples were collected prior to commencement of treatment and after completion of neoadjuvant therapy prior to surgical resection. In the Cambridge cohort, additional blood samples were collected at 6 weeks and 12 weeks during neoadjuvant treatment.

### DNA extraction from tumor and germline samples

For the Mayo cohort, tumor DNA was extracted from four 10 micron sections obtained from archived formalin-fixed paraffin-embedded tissue using the MagMAX FFPE DNA/RNA Ultra Kit (ThermoFisher Scientific), following macro-dissection to enrich for tumor cells guided by an H&E stained tumor section. For the Cambridge cohort, tumor DNA was extracted from ten 30μm sections obtained from the fresh frozen tumor tissue using the DNeasy Blood and Tissue Kit (Qiagen). Germline DNA was extracted from peripheral blood cells using the DNeasy Blood and Tissue Kit (Qiagen).

### Plasma processing, DNA extraction and quality assessment

Blood was collected in 10 mL K2 EDTA tubes and centrifuged at 820g for 10 minutes within 3 hours of venipuncture to separate plasma. 1 mL aliquots of plasma were centrifuged a second time at 16000g for 10 minutes to pellet any remaining leukocytes and the supernatant plasma was stored at −80 °C. cfDNA was extracted using either the QIAsymphony DSP Circulating DNA Kit (Qiagen) or MagMAX Cell-Free DNA Isolation kit (ThermoFisher Scientific). All cfDNA samples were evaluated for yield and quality using droplet digital PCR, as described previously(*21*).

### Tumor/Germline Exome Sequencing

For the Mayo cohort, tumor/germline exome sequencing libraries were prepared using 44-200 ng of tumor DNA (mean 175 ng) and 148-200 ng of germline DNA (mean 195 ng), using the KAPA Hyper Prep Kit following manufacturer’s instructions. Exome enrichment through hybridization was performed using a customized version of Agilent SureSelect V6 exome. For the Cambridge cohort, tumor and germline exome libraries were generated using the Illumina Nextera Rapid Capture Exome Library Preparation kit, using 50ng of DNA as input. We pooled exome libraries and sequenced on Illumina HiSeq.

### Variant calling in tumor exomes and identification of target mutations

Reads were aligned to human genome version hg19 using bwamem(*22*), followed by base recalibration using GATK(*23*), duplicate identification using Picard tools MarkDuplicates, and indel realignment using GATK. Germline mutations were inferred using GATK HaplotypeCaller and Freebayes(*24*). Somatic tumor mutations were called using MuTect(*25*), Seurat(*26*) and Strelka(*27*). Somatic mutations with an allele frequency < 5% were removed.

### Identification of Target Mutations

Potential target mutations found on autosomal chromosomes were assessed for copy number, purity, and variant allele frequency (VAF). We used Sequenza to infer both the proportion of tumor cells in the sequenced tumor DNA sample and copy number alterations in the tumor(*28*). For each mutation, the mean variant allele frequency from the variant callers, sample purity, and local copy number were used to infer its cancer cell fraction (CCF) via two different methods: an implementation of the algorithm from McGranahan et. al.(*29*), and PyClone(*30*). For each sample, the VAF, minor and major copy number, and purity were used as input for PyClone analysis with 25,000 iterations, including 10,000 iterations of burn in.

Founder mutations were identified using a set of criteria for mutation confidence and maximum CCF. To quality as a target for ctDNA analysis, a mutation must have been identified by at least 2 somatic mutation callers, have a mean of >20x germline reads passing each mutation caller’s filters that covered the mutated base, a germline VAF < 0.01%, and >50x mean tumor passing filter reads. In addition, the upper range of the CCF distribution calculated using the McGranahan et. al. approach must be equal to 1.0, and the mutation must be found in the highest CCF PyClone mutation cluster.

### Primer Design for TARDIS

Mutations that passed the filtering steps above, plus any mutations found in genes of interest(*31*), were used as targets for TARDIS primer design. The primer design process is focused on maximizing TARDIS performance and minimizing spurious amplification, particularly in the linear preamplification stage. We first generated primers on the forward or reverse strands up to 350 bp from the target mutation position for both linear (preamp) and exponential (PCR1) amplification reactions using Primer3(*32*). Preamp primer melting temperature (Tm) range was set to 68-74 °C, and PCR1 Tm range was 56-60 °C, with the Preamp primer upstream and a maximum of 3bp overlap allowed between Preamp and PCR primers. We also require the 3’ end of the PCR1 primer to be between 3bp and 10bp from the target mutation position, to ensure short mutant molecules are captured efficiently but the targeted mutation is not masked by potential primer synthesis overhangs. To avoid unintended amplification in multiplexed PCR reactions, we used a combination of in silico PCR, sequence comparison to the genome using LAST(*33*), and 3’ primer kmer matching to identify problematic primers for multiplexing. Each primer for each target is evaluated for kmer matches to the regions around other target mutations, and the primers with the fewest number of matches to other target regions were selected for each target. We then constructed a network with each node representing a target/primer and edges representing 6mer matches between the 3’ end of a target’s preamp primer and the sequence 150bp on either side of another target, or in-silico PCR predicted amplicons generated by the two preamp primers. Finally, we iteratively removed the node with the most edges until there were no remaining edges. If there were target mutations that must be included in a panel, such as driver mutations, all primers with 3’ edges to these targets were removed first to ensure they remain in the final target set. A test run of TARDIS using each new primer panel was conducted with 8 replicates of sheared genomic DNA before analyzing plasma samples to identify any remaining problematic primers. An amplicon was removed from the panel prior to analysis of plasma samples if median proportion > 0.5 or maximum proportion >0.75 of the reads for that amplicon were masked or if the amplicon captured a median proportion >0.5 or maximum proportion >0.75 of all reads in any control run.

### Preparation of TARDIS sequencing libraries

4.0-19.0 ng of template plasma DNA (mean 10.4, median 10.3) for linear pre-amplification (preamp). For each TARDIS run, patient-specific primers were pooled equimolarly. Each pool was used at a final concentration of 0.5 *µ*M (regardless of the number of primers in the panel). Amplification was performed using Kapa Hifi HotStart Mastermix (Kapa Biosystems) at the following thermocycling conditions: 95°C for 5 minutes followed by 50 cycles at 98°C for 20 seconds, 70°C for 15 seconds, 72°C for 15 seconds, and 72°C for 1 minute. This reaction was followed by a magnetic bead cleanup (SPRIselect, Beckman Coulter) at 1.8x ratio after addition of 10% ethanol. Preamplified DNA was eluted in 10 *µ*L water. After dephosphorylation using FastAP (ThermoFisher Scientific), 0.8 *µ*L of 100 *µ*M ligation adapter was added to each sample. The sequence of the hairpin oligonucleotide using for single-stranded DNA ligation is provided in Table S4 and was adapted from Kwok et al.(*34*). Samples were denatured at 95°C for 5 minutes and immediately transferred to an ice bath for at least 2 minutes. We setup ligation reactions using 2.5 *µ*L 10x T4 DNA Ligase buffer (New England Biolabs), 2.5 *µ*L of 5 M betaine, 2,000 U of T4 DNA ligase (New England Biolabs) and 5.8 *µ*L of 60% PEG8000. Ligation was performed at 16 °C for 14-17 hours. A magnetic bead cleanup (SPRIselect) was performed at 1x buffer ratio after initially diluting the sample by adding 40 µL water (to reduce effective PEG concentration during cleanup). An additional dephosphorylation was performed using FastAP.

Exponential PCR (PCR1) was performed using nested primers downstream of the pre-amplification primers and a universal reverse primer complementary to the ligated adapter, upstream of the UMI (see Table S4 for primer sequences). Primers were pooled equimolarly and used at a final pool concentration of 0.5*µ*M. We used the NEBNext Q5 Hot Start HiFi PCR Mastermix (New England Biolabs) with the following thermocycling conditions: 98°C for 1 minute followed by 5 cycles at 98° for 10 seconds, 61.5°C for 4 minutes, and 15 cycles at 98° for 10 seconds, 61.5°C for 30 seconds and 72°C for 20 seconds, followed by a 2 minute incubation at 72°C. After a 1.7x magnetic bead cleanup (SPRIselect), we eluted the product in 40 *µ*L water. A second round of PCR (PCR2) was performed using universal primers to introduce sample specific barcodes and complete sequencing adaptors, as described previously(*12*). We used 1 U per reaction of Platinum Taq DNA Polymerase High Fidelity (Invitrogen) in the following buffer: 1.3x Platinum buffer, 0.4M betaine, 2.5 *µ*l/rxn of DMSO, 0.45mM dNTPs, 1.75 mM MgSO_4_ and primers at 0.5 *µ*M. 10 μL of the product from the first amplification was used as template, at the following thermocycling conditions: 94°C for 2 minutes followed by 15 cycles at 94°C for 30 seconds, 56° for 30 seconds, 68°C for 1 minute, and a final incubation at 68°C for 10 minutes. A final magnetic bead cleanup (SPRIselect) was performed at 1.2x volume ratio. TARDIS libraries were eluted in 20 µL DNA suspension buffer, quantified using qPCR (KAPA SYBR FAST Universal qPCR kit, Kapa Biosystems) and pooled for sequencing. Sequencing was performed on Illumina HiSeq 4000 or Illumina NextSeq.

### Analysis of TARDIS Sequencing Data

Paired-end sequencing reads were aligned to human genome hg19 using bwa-mem. Read pairs whose R1 read mapped to the start position of a target primer were considered on-target reads, while the position of the R2 read was used to determine the length of the template molecule. The UMI sequence and molecule size were used to identify all of the reads that came from the same template molecule. To minimize incorrect assignment of reads to read families, we implemented a directed adjacency graph approach inspired by Smith et al.(*35*). Briefly, a graph is constructed in which each UMI is a node. An edge from nodeA to nodeB is created if their UMIs differ by one base, their DNA molecule size is the same, and nodeA has at least twice as many reads as nodeB. All of the reads from UMIs in each component from the resulting graph constitute a read family and are considered to have come from the same original molecule. UMI variation within a read family is assumed to arise due to PCR or sequencing error. We found that a small number of UMIs with very few reads had incoming edges from multiple otherwise separate components. The component assignment of these nodes is ambiguous, and they significantly reduced the number of independent components in the graph. To resolve this issue, any UMI that had two or more incoming edges and no outgoing edges was removed. We then inferred the allele at the target position by consensus of all R1 reads in a given component, requiring that at least 90% of the R1 reads carried a particular allele at the position of interest. In practice, the vast majority of read families contained fewer than 10 reads, and therefore required perfect agreement at the target position. Inferred molecules with less than 90% read support for a variant were removed as inconclusive.

Allele fractions (AFs) were calculated as either number of mutant reads as a fraction of total reads (Raw AF) or number of mutant RFs as a fraction of total RFs (TARDIS AF). To ascertain ctDNA detection in a sample, we required support of at least 2 RFs. For any mutations supporting ctDNA detection, we required that its TARDIS AF represent at least 0.5 mutant molecules in the reaction. In addition, the ratio between number of RFs supporting a mutation and mixed RFs observed at that locus must be <15. Finally, if only mutation supported ctDNA detection, we required at least 2 independent RF sizes (to ensure independent PCR events). This requirement was waived if >1 mutation supported ctDNA detection. To quantify ctDNA levels, we calculated mean AFs over all targeted mutations using RawVAFs – to avoid limiting the quantification to only DNA fragments that are successfully represented by an RF. However, to avoid the contribution of background noise, RawVAFs for any mutations not supported by ≥1 RF, a ratio with Mixed RFs of ≥15 or <0.5 mutant molecules were set to zero prior to calculating the mean.

### TARDIS Analysis Pipelines

Target selection and primer design pipelines were developed in Python3 using NumPy, SciPy, networkX, pandas, and matplotlib, and in Julia 0.6.2 using BioJulia, DataFrames, Gadfly, and LightGraphs. Data analysis and plotting were conducted in Python3, Julia 0.6.2, and R v3 using ggplot2.

### Calculation of background error rates

To measure overall background error rates, we evaluated the first 10 bp at each locus for highest non-reference alleles (starting 2 bp downstream of target-specific primers). For 8 target amplicons across 6 sheared DNA control replicates, our evaluation dataset contained a total of 480 independent positions covering 80 genomic loci. In raw sequencing results, we observed an error rate of 4.68 × 10^−4^, with background errors observed at all loci. We found that requiring consensus of all members of an RF significantly reduced error rates (Fig. S2). We observed 15/80 amplicon positions with 181 unexpected variants using RF consensus. 84 of them (46.4%) were contributed by 5/80 (6.25%) genomic loci, suggesting either detection of low abundance alleles present in the DNA sample (obtained from cell lines) or that these loci are highly error prone. Excluding these 5 loci, we observed a mean error rate of 1.03 × 10^−4^, contributed by 10/75 representative loci. By requiring each putative error to be supported by RFs of at least 2 sizes, we observed 98 unexpected variants, of which 70 (71.4%) were contributed by the 5 error prone loci. Removing these 5 loci, our mean error rate was 3.29 × 10^−5^ contributed by 8/75 representative loci while the remaining were error free in all replicates.

## Acknowledgments

We thank Marissa Pacheco and Leslie Dixon at Mayo Clinic for their support in collection and processing of patient samples. We thank all the patients for their participation in this study.

## Author contributions

BAP and MM conceptualized and designed the study. BRM, TCC, NP and MM developed methods. SJS, BE, PAC, KSA, HEK, DWN, AEM, BKP, CC and BAP designed and conducted the prospective clinical studies. TCC, SJS, AOB and MF generated data. BRM and MM analyzed sequencing data. BRM, SJS, CC, BAP and MM interpreted data. BRM and MM wrote the paper the assistance from SJS, AOB, CC, BAP and other authors. All authors approved the final manuscript;

## Data and materials availability

Tumor/germline sequencing data and targeted sequencing data from plasma analysis will be deposited in a public database and made available at manuscript publication.

